# Huddling nest use are the optimal strategies for heat conservation in a social marsupial: lessons from biophysical models

**DOI:** 10.1101/2022.05.29.493884

**Authors:** Roberto F. Nespolo, Isabella Peña, Carlos Mejías, Abel Ñunque, Tomás Altamirano, Francisco Bozinovic

## Abstract

Endothermy, understood as the maintenance of continuous and high body temperatures (T_B_) due to the combination of metabolic heat production and an insulative cover, is severely challenged in small endotherms of cold environments. As a response, social clustering, and nest use (collectively, “communal nesting”) are common strategies for heat conservation in small mammals and birds. To quantify the actual amount of energy that is saved by communal nesting, we studied the social marsupial *Dromiciops gliroides* (monito del monte), a relict marsupial species from the cold forests of southern South America. It is hypothesized that sociability in this marsupial was driven by the cold, for which we calculated the energetic benefits that communal nesting confers. Using biophysical models and experimental coolings, we simulated heat exchanges experienced by grouped or solitary individuals, and also individuals within nests, collected from the field. Assuming a model of passive cooling, we calculated the net energy cost of euthermic maintenance (E_cost_: the total energy needed to maintain euthermia). We adjusted 50 cooling curves, to exponential decay models, and found in all cases that the strategy minimizing heat loss is to be clustered within a nest, for which the E_cost_ was the minimum. This was significantly lower than the clustered condition, outside the nest, a reduction that represents almost half of energy consumption per day in a resting, thermoneutral condition for this marsupial. Overall, our results suggest that the strategy that significantly maximized heat conservation, compared with alternative strategies, was communal nesting. These findings support the idea that, in this social mammal, sociality is driven by bioenergetic benefits.

## Introduction

Organisms exist in thermodynamic equilibria with their environment (Porter and Gates, 1969), and cold conditions create especially tough challenges for small, endothermic animals with high rates of heat dissipation, due to their high surface-to-volume ratio (Canals et al., 1989). Although fur and feathers provide good insulation, at very small sizes these are not enough to maintain sustained euthermia. Thus, other strategies were promoted by natural selection, some are physiological, such as increase in thermogenic capacity and heterothermic responses (Bozinovic et al., 1987; Nespolo et al., 1999; Sharbaugh, 2001); while some others are behavioral, such as the formation of clusters or huddling and the use of nests (=communal nesting) (Bozinovic et al., 1988; Canals et al., 1989; Canals et al., 1997; Gilbert et al., 2010; Scholander, 1955).

The fact that clustered individuals conserve heat more efficiently than isolated ones, is a generalized biophysical consequence of the exponential reduction of surface-to-volume ratio (Canals et al., 1997). That’s why huddling is so widespread in small birds and mammals of cold environments (e.g., marsupials: Fisher et al., 2011, rodents: Antinuchi and Busch, 2001; Kauffman et al., 2003, birds: Lubbe et al., 2018; see reviews in Gilbert et al., 2010; Wojciechowski et al., 2011). However, communal nesting is also attributed to other factors, such as social behavior, protection and structural support (reviewed by Ebensperger and Labra, 2020). For instance, in the Siberian flying squirrel clustering behavior is explained by subsequent mating rather than kingship or thermoregulating benefits (Selonen et al., 2014). The avian cup-shaped nest design is primarily explained by structural support, and not insulation (Heenan and Seymour, 2011). Similarly, in the social “degu” (*Octodon degus*), communal nesting is explained by kinship, rather than thermoregulation (Ebensperger and Bozinovic, 2000; Ebensperger et al., 2004). However, it has been challenging to test these competing hypotheses on live animals, either in the laboratory or in the field, due to the obvious limitations of obtaining the animals, the precision of the technique for measuring energy consumption, or even obtaining the nests.

Here we took advantage of a collaborative effort focused on the social marsupial monito del monte (*Dromiciops gliroides*), which normally cluster together and build sophisticated nests. *D. gliroides* is an arboreal small mammal (∼30g) endemic to the temperate rainforests of southern South America, which includes high Andean locations where winter temperatures reach freezing values, but also mild locations near the coast with mean winter temperatures of about 5ºC (Gurovich et al., 2015; Mejias et al., 2021). Monitos elaborate complex nests, built by interlaced quila leaves (*Chusquea quila*, an endemic bamboo), covered with moss inside, and the whole structure is usually located within tree cavities (Celis-Diez et al., 2012; Franco et al., 2011; Honorato et al., 2016). The nests are spherical, with a single entrance, built with plant materials from *Chusquea* spp. leaves, *Hymenophyllum* spp. ferns, and lined with ferns and mosses, which provide both thermal insulation and structural support (Honorato et al., 2016; Vazquez et al., 2020).

It is likely that monitos do communal nesting as a strategy to cope with the cold, but two facts complicate this conclusion. *First*, direct measurements of energy expenditure in clustered and isolated individuals did not produce significant differences between treatments (Franco et al., 2012). *Second*, monitos are one of the few marsupials with advanced levels of sociality (Fonturbel et al., 2022; Nespolo et al., 2022b; Russell, 1984). The sociality of monitos is supported by observations of long-term persistence of familiar groups and the existence of foraging groups composed of unrelated individuals, in spring and summer (Fonturbel et al., 2022; Nespolo et al., 2022b).

Here we simulated clustering, combined with nest use, and testes if these strategies are synergistic in conferring energetic savings in term of heat conservation. If communal nesting represents an adaptive strategy to cope with the cold, then clustered individuals within nests should conserve the heat significantly longer than isolated individuals without nests.

## Material and methods

We used real monitos nests, that were obtained from a bird conservation program in charge of one of the coauthors (T. Altamirano). We had access to 240 nestling boxes located at Pucón (39º18’51’’S; 71º52’50’’W, 1100 m.a.s.l.), that are normally colonized by birds in spring, and monitos in autumn. Each nesting box (40×30×30cm, wood) was placed hanging from the trees about 1.5 m above the ground and separated 30-40m from each other (Altamirano et al., 2015). During the 2019 winter, we obtained 17 monitos nests (dry weight 55.1± 2.4 g, mean±se) (Fig 1), that were found within the nest box at the end of the winter. Each nest was individually placed in paper bags and stored at 4ºC until our experiments, two months later.

**Fig 1a).**
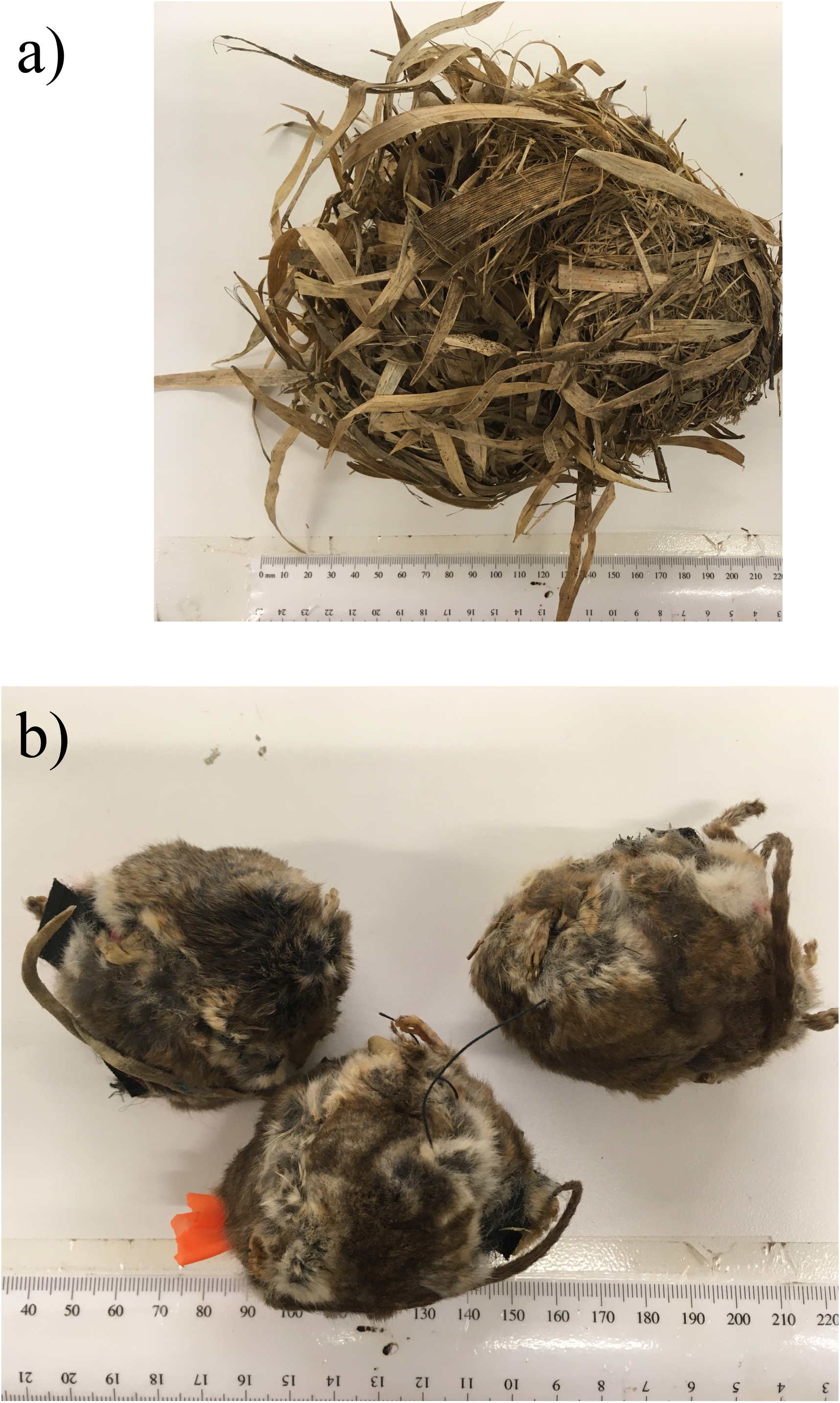
Monitos nests used in this study; b) the mannequins used for the cooling curves.

The taxidermic models (“mannequins”, hereafter) were manufactured with skins of dead monitos, provided by the National Services for Wildlife (SAG and CONAF). Nine carcasses were used, which were fresh-dissected and left for 2-3 days stretched, covered by a layer of sodium chloride and boron sulphate to dry out. Two skins were needed to manufacture one individual, thus we built five mannequins in total (Fig 1). Each mannequin was fabricated using a 40g agar bead, which was covered by the skin and sutured with surgical sutures. In the middle of the agar sphere, a 4(diameter)x9(length) mm cylindrical space was left to introduce the temperature sensor (see below), and a Velcro closure was included for opening. The agar was prepared with 3 grams of agar-agar and 100 ml of boiling water. The sphere was wrapped in a rubber balloon so that the agar did not dehydrate at high temperatures. The mannequin’s weight was: 55.1 ± 2.4 g, with a volume of 39.8 ± 2.7 ml, and a density of 1.4 ± 0.09 g/ml.

For the cooling curves, we used two “Pelt” chambers (60×40×30cm, Sable Systems, USA) to maintain constant temperatures, one for warming up the mannequins to 40 ° C ± 1 ° C and the other for cooling it at 5 ° C ± 1 ° C (the actual cooling curve). We located a wire frame in the floor of the chamber so that neither mannequins nor nests were in direct contact with the chamber material. For each cooling curve, the procedure consisted of removing the mannequin from the warm chamber, waiting for it to cool down to 35ºC (the euthermic body temperature of a monito), and placing it within the cold chamber. We performed two cooling curves per day. To continuously record the core temperature of the mannequins, we used temperature data loggers, (model DST-micro, Star Oddi, Iceland; cylindrical 25 millimeters long and 5 mm in diameter), with a data-gathering frequency of one per minute. According to the manufacturer, the devices are calibrated at factory over a temperature range of 5 to 45ºC. We also tested the devices in a beaker with water at 40ºC that was allowed to cool to room temperature (10ºC), with temperature records every two minutes, using a copper-constantan thermocouple (Cole Parmer). The linear regression between water and logger temperature (20 points) was highly significant (R^2^ = 0.99, p=0.001). The environmental temperature within the chambers was recorded, in addition to the chamber’s thermocouple, with a HOBO environmental temperature data logger, model pro v2 (USA) attached to the internal walls of the chamber, with the probe hanging to the air. This probe was configured to record one data per minute. Every cooling had a duration of 593 minutes, and we performed a total of 50 curves combining solitary, clustered and individuals within nests. That is, we ran five curves with bare datalogger, five curves with the agar sphere (no fur), 10 curves with a solitary mannequin, 10 curves with a solitary mannequin within a nest, 10 curves with a group of 3 mannequins and 10 curves with a group of 3 mannequins within a nest. For visualization of heat distribution, we took IR thermographic images at different stages of the warming, using a thermograph (FLIR i40, Sweden).

Each cooling curve was adjusted to a one-phase exponential decay given by the equation: y = (y_0_-plateau)e^-k*time^+plateau, where y_0_ is y at time=zero, the plateau is the asymptotic temperature, and k is the rate constant, expressed as the reciprocal of time (the higher the ks, the higher the heat loss rate). We also calculated the time constant (tau), as the reciprocal of k, and the half-time, as ln(2)/k.

In addition to estimating curve parameters, we predicted the euthermic cost of maintenance by calculating, for every minute, the energy that a monito would spend to maintain its body temperature constant (E_cost_). Thus, we used the Newton’s cooling equation for passive cooling, which is normally used for estimating metabolic rates in steady-state conditions among endotherms (Mejias et al., 2022; Nicol and Andersen, 2007; Rezende and Bacigalupe, 2015), and calculated E_cost_ as: C(T_B_-T_A_), where C is minimum thermal conductance of euthermic *D. gliroides*, (C=3.4848 Jg^-1^h^-1^ºC^-1^)(Bozinovic et al., 2004). Then, we compared the overall E_cost_(=net E_cost_) for the entire cooling curve of, among the different conditions. Both, curve parameters and net E_cost_ were compared among the six treatments: bare data-logger, data-logger with agar, data-logger with fur (=mannequin), grouped mannequins (3 individuals), individual mannequin within a nest and grouped mannequins within a nest. These were compared with a one-way ANOVA, using the Statistica software (Statistica, 2006). Parametric assumptions were checked by the Levene test for homogeneity of variance and Shapiro-Wilks for normality.

## Results

The IR thermal images showed how the core of the body maintained the heat (33.8ºC, in red) until the fourth hour of the cooling, while maintaining a cooler envelope (skin with fur, in yellow, Fig 2a-d). These images also show, qualitatively, the better heat conservation of clustered individuals compared with solitary ones. The adjustment of every one of the 50 cooling curves had an R^2^ above 0.98 (all curves are plotted in Fig 3).

**Fig 2.**
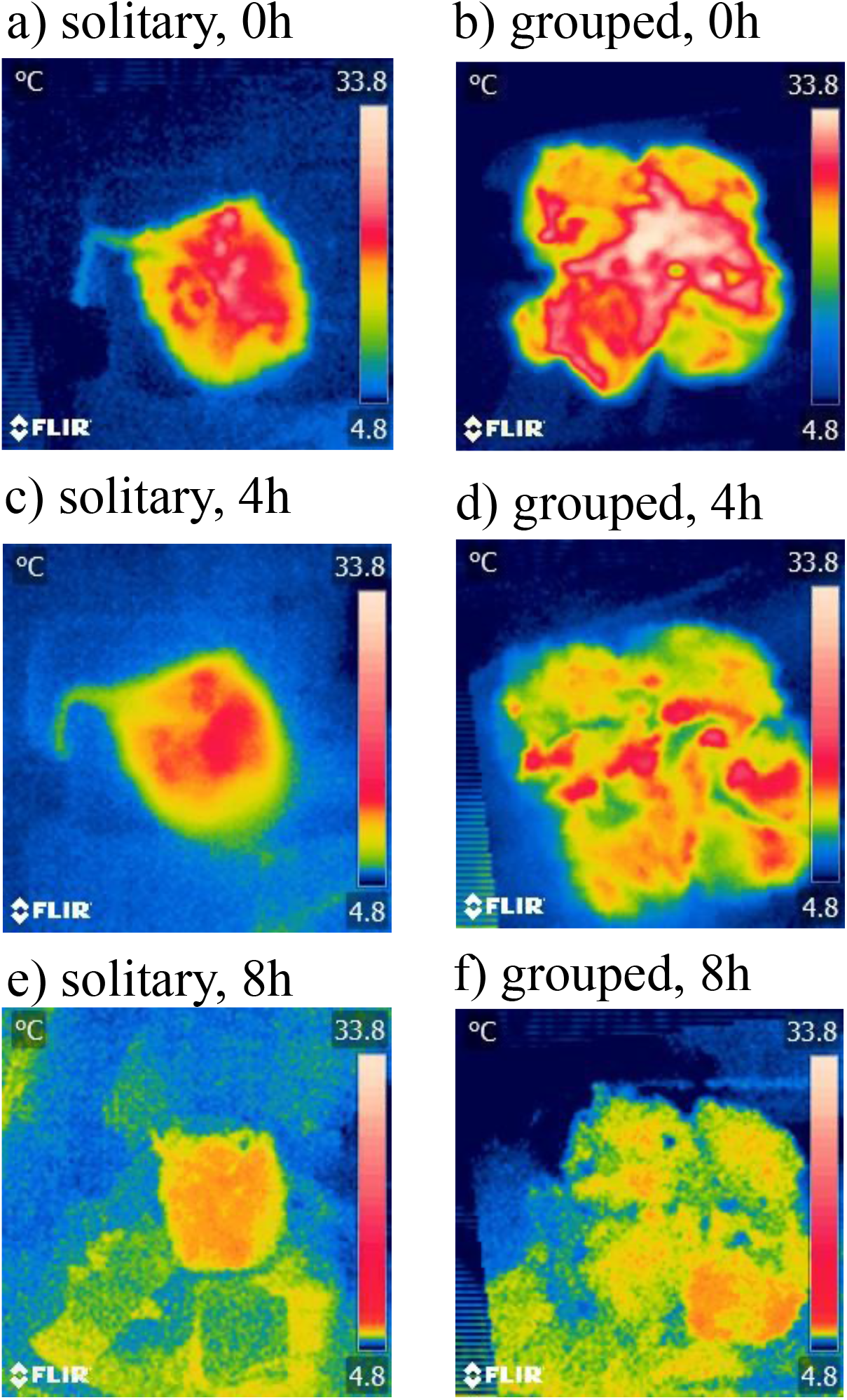
Representative thermographic images obtained from grouped and solitary mannequins at different moments of the cooling. The images show how the outermost surface of the skin with fur is colder than the core, thus acting as an isolating envelope.

**Fig 3.**
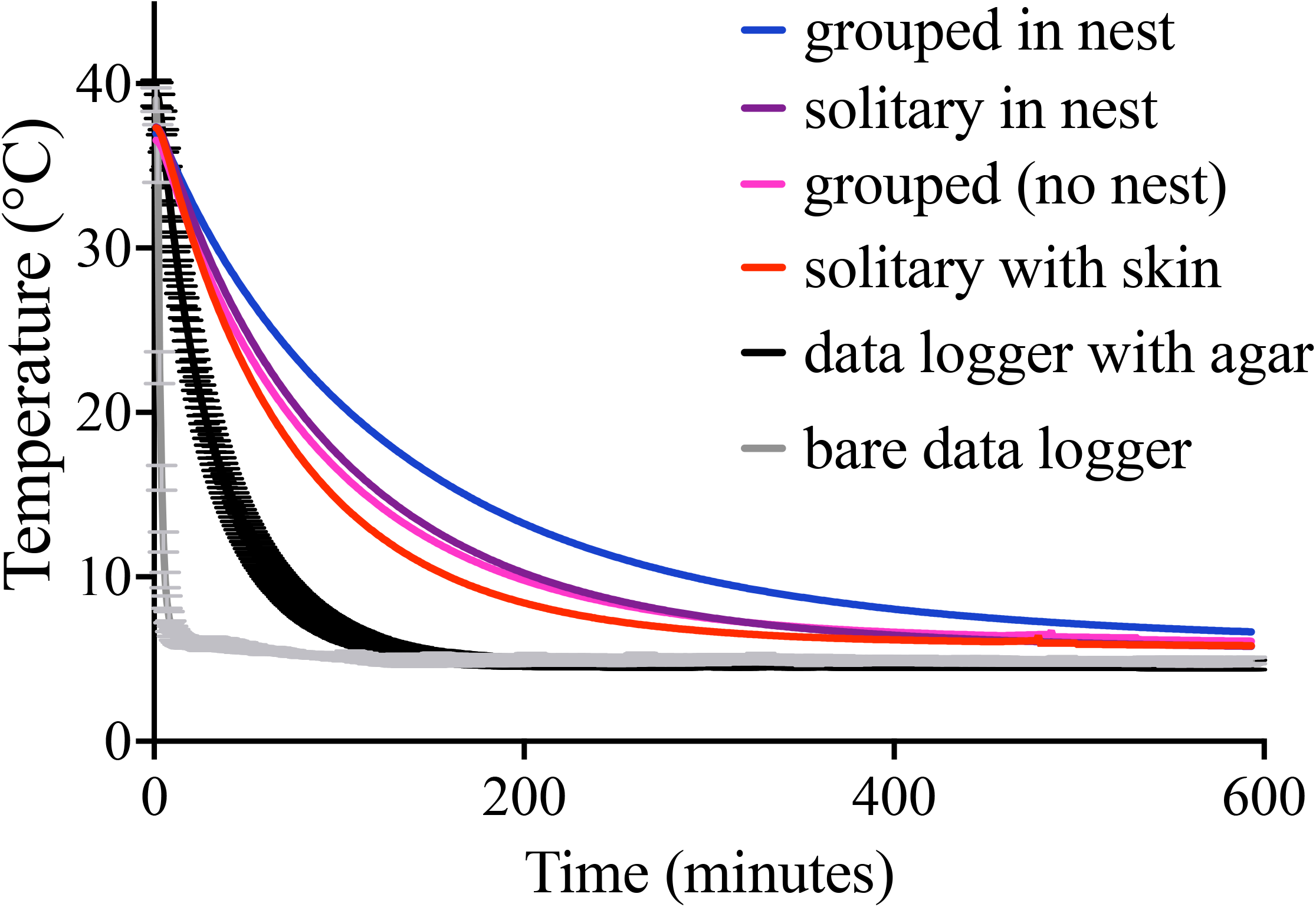
Cooling curves and experimental treatments. All curves are plotted.

We found significant differences among treatments in rate constant (F_5,43_=495.5, P=0.0001, one way ANOVA), half time (F_5,43_=65.7, P=0.0001, one way ANOVA) and time constant (F_5,43_=65.8, P=0.0001, one way ANOVA). The agar (i.e., the difference in cooling rate of the data logger with and without agar) reduced the rate of heat loss 12 times (from 0.37±0.02 to 0.031±0.002 min^-1^, P=0.001, Fisher post-hoc test, Table 1). On the other hand, the fur (i.e., the difference in cooling rate of the mannequin with and without skin) reduced the rate of heat loss 2.3 times (from 0.031±0.002 to 0.014±0.0006 min^-1^, P=0.001, Fisher post-hoc test, Table 1). However, the rate constant was non-significantly different between solitary, grouped, solitary in nests and grouped in nests mannequins (P>0.Fisher post-hoc tests, Table 1). Given that half-time and time constant are both measures of cooling speed, we will interpret half-time. Grouped mannequins within nests conserved heat 31% longer (half-time: 91.7 ± 4.71 min) than solitary mannequins in nests (half-time: 63.2 ± 2.8 min; P=0.008, Fisher post-hoc test, Table 1). Solitary individuals, however, conserved heat 1% lower (non-significantly different) than grouped individuals without nests (half-time: 64.1 ± 3.6 min; P= 0.43, Fisher post-hoc test, Table 1). Solitary individual in nests in turn, conserved heat 18% higher than solitary individuals without nest (half-time: 64.1 ± 3.6 min versus 52.5 ± 3.0 min), which was also significant (P= 0.021, Fisher post-hoc test, Table 1).

**Table 1.** Mean (± standard errors) of the exponential decay parameters of cooling curves (n=5 for data logger and data logger with agar, and n=10 for the rest). Different letters in boldface represent statistically significant differences (P<0.01) after a one-way ANOVA and a Fisher post-hoc test.

| Treatment | rate constant<br>(min<sup>-1</sup>) |  | half-time<br>(min) |  | time constant<br>(min) |  |
| --- | --- | --- | --- | --- | --- | --- |
| data logger | 0.367**a** | $\pm 0.02$ | 1.9**a** | 0.1 | 2.8**a** | 0.2 |
| data logger + agar | 0.031**b** | $\pm 0.002$ | 23.0**b** | 1.7 | 33.2**b** | 2.5 |
| data logger + agar + fur<br>(solitary individual) | 0.014**c** | $\pm 0.0006$ | 52.6**c** | 3.0 | 75.8**c** | 4.4 |
| grouped (n=3) | 0.011**c** | $\pm 0.0006$ | 64.1**d** | 3.6 | 92.5**d** | 5.1 |
| solitary in nest | 0.011**c** | $\pm 0.0005$ | 63.2**d** | 2.8 | 91.2**d** | 4.0 |
| grouped in nest | 0.008**c** | $\pm 0.0004$ | 91.7**d** | 4.7 | 132.3**e** | 6.8 |

The difference in half-time between mannequins with fur and without fur was highly significant (56%) (52.6 ± 3.0 min versus 23.0 ± 1.7 min, P= 0.0001, Fisher post-hoc test), and also the difference between the bare datalogger and the datalogger with agar (23.0 ± 1.7 min versus 1.9 ± 0.11 min, P= 0.0001, Fisher post-hoc test, Table 1). Hence, in terms of heat conservation, grouped individuals outside nests are equivalent to solitary individuals within nests (Table 1). The amount of metabolic heat that a monito should produce every minute, to maintain constant body temperature represents the euthermic cost of maintenance (E_cost_). This was calculated using the Newton’s passive cooling equation (see Methods) and estimated for clustering mannequins (mimicking a group of monitos as shown in Fig 4a) and mannequins within nests (Fig 4b), and presented in Fig 4c. This comparison shows that the configuration that minimizes E_cost_ is to be grouped within a nest (red line in Fig 4c). Summing E^costs^ across the cooling period confirms that the net euthermic cost of maintenance is minimized by the combined use of nest in groups (Fig 4d). Indeed, significant differences were found after a one-way ANOVA (F_5,44_=49.4, P=0.0001, Fig 4d). Specifically, to be grouped in the nest has an E_cost_(44.5 ± 0.83 kJ) that is significantly lower than to be grouped outside a nest (48.1±1.38 kJ, P=0.008, Fisher post-hoc test). However, to be solitary in the nest (50.8±0.62 kJ) is equivalent (not significantly different) with to be grouped without nest (48.1±1.38 kJ, P=0.07, Fisher post-hoc test). There were significant differences in E_cost_ between the mannequin with and without fur (51.4±0.55 versus 62.1±1.85 kJ, P=0.0001, Fisher post-hoc test). Finally, the calculated E_cost_ of the bare data-logger and the mannequin without fur was non-significant (64.2±1.70 versus 62.1±1.85 kJ, P=0.26, Fisher post-hoc test).

**Fig 4.**
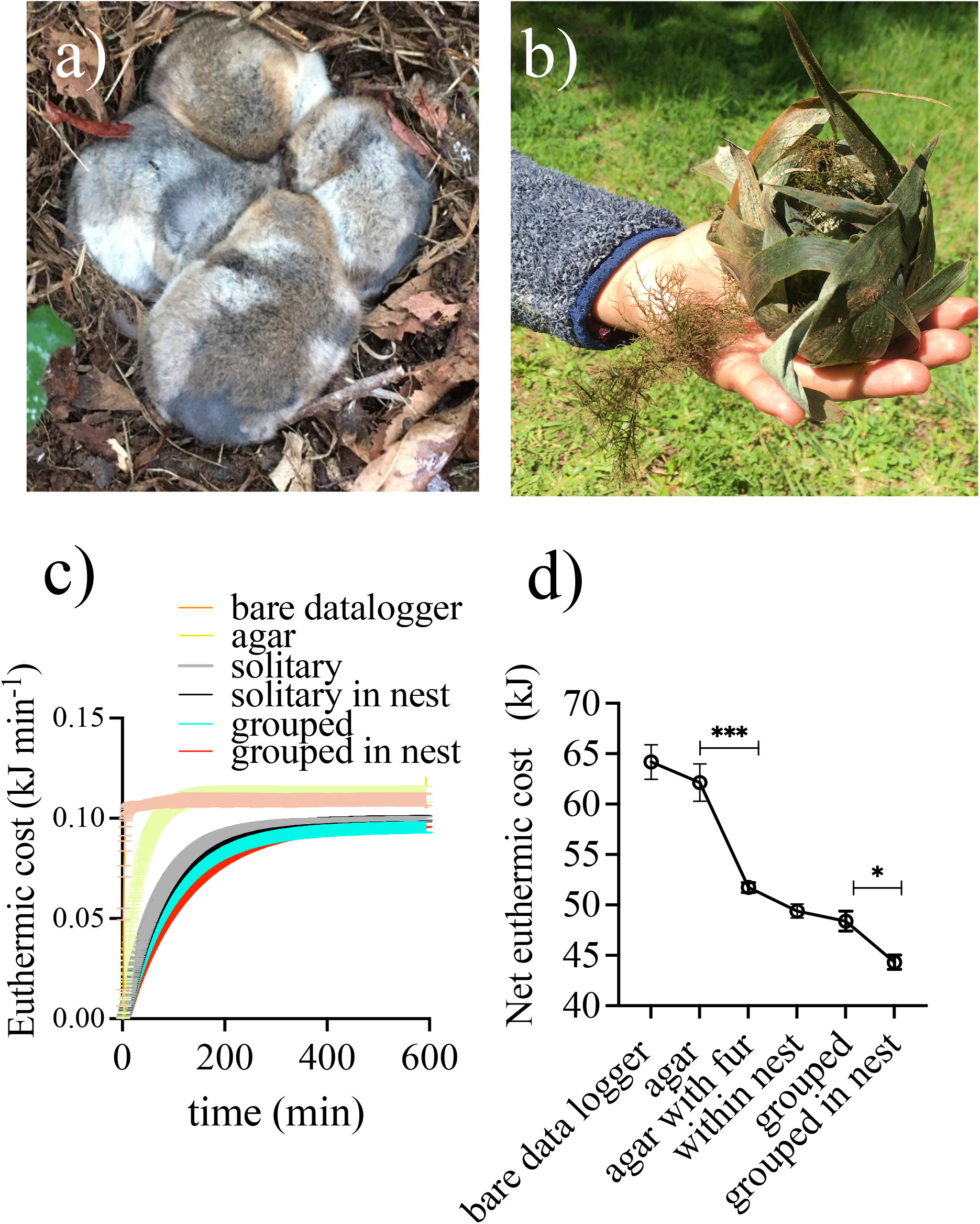
a) clustered (live) individuals in a typical configuration found in tree holes and nests (photo: R.Nespolo); b) a “fresh” nest found after 2020 winter; c) euthermic costs of maintenance per minute, calculated during experimental cooling from 35 to 5ºC in a climatic chamber (see methods for details). This calculation assumes that the animal replaces the heat loss by an equivalent amount of metabolic heat production (Newton’s passive cooling); d) the net euthermic cost of maintenance (mean ± se) over the complete cooling period. Asterisks denote significant differences between groups after a one-way ANOVA and Fisher post-hoc test (*p<0.05; **p<0.01; ***p<0.001).

## Discussion

### Communal nesting, bioenergetics and sociality

There is an historic discussion about whether nest sharing in clusters has a bioenergetic significance (i.e., a heat-conservation strategy) or represents a by-product of the benefits of group living (Dausmann and Glos, 2015; Ebensperger, 2001; Franco et al., 2011; Gilbert et al., 2010; Heenan and Seymour, 2011; Lubbe et al., 2018; Madikiza and San, 2020; Olson et al., 2018; Selonen et al., 2014; Vickery and Millar, 1984; Vogt and Lynch, 1982; Willis and Brigham, 2007; Withers and Jarvis, 1980). The problem arises because often, species with advanced levels of sociality also benefit from clustering during cold periods. For instance, in the dormouse (*Glis glis*) social clustering seems to be explained by mating behavior rather than by thermoregulatory benefits (Madikiza and San, 2020), and in North American flying squirrels (genus *Glaucomys*), interspecific social nestling appear to mainly driven by their sociality (Olson et al., 2018; Selonen et al., 2014). However, in birds, it has been documented that social thermoregulation combined with the use of insulative roosts and nests, provide energy savings over 50%, compared with isolated birds outside nests (Lubbe et al., 2018). This is also the case of Siberian hamsters, which replace experimental shaving by having access to nest materials (Kauffman et al., 2003). Nest use and clustering, however, confer slight amelioration of cold stress in laboratory mice (Lubbe et al., 2018). Another important effect of the use of nests and clustering is the reduction of lower limit of thermoneutrality, and remain more time at the BMR (Bozinovic et al., 1988). Our results, suggesting that the optimal strategy is clustering and nest use, is in consonance with thermodynamic predictions.

### Impact on energy budget

In the field, monitos exhibit high phenotypic flexibility in nesting building behavior, sometimes building nests within tree cavities and sometimes outside them (Vazquez et al., 2020). Also, both the mass and the volume of the nests increase with elevation, thus suggesting that animals build more insulated nests in colder environments (Altamirano et al., 2019). This strategy is combined with huddling behavior, as larger groups are often found in colder locations (Altamirano et al., 2019; Celis-Diez et al., 2012). Indeed, according to (Canals et al., 1997), the average relative area lost during huddling in small mammals ranges from 28.7 to 39.1%, where the maximum reduction in surface area is attained with three individuals. Indeed, according to Bozinovic et al. (1988), the reduction in minimum thermal conductance due to huddling and nest use is about 42% (compared with animals housed individually), which is a significant fraction of the energy budget of a small marsupial or rodent, especially in winter. However, these authors used n=5 individuals and experimental temperatures of -10ºC, which is outside the natural range for the case being discussed. In our experiments (with minimum T_A_ of 5ºC), the nests provided additional energy savings of about 10% (a reduction in E_cost_ 49 to 44 kJ, see Fig 4d). Considering that the basal metabolic rate (BMR) of *D. gliroides* is 13 kJ d^-1^, these 5kJ represents almost half of energy consumption per day in a resting, thermoneutral condition (Nespolo et al., 2022a).

### Communal nesting and hibernation

In heterothermic species (i.e., animals that express daily or seasonal torpor, *sensu* (Ruf and Geiser, 2015), for which *D. gliroides* is an example (Nespolo et al., 2021), huddling has been associated to increased survival (Boratynski et al., 2015; Boyles et al., 2008; Patil et al., 2013). However, when animals experience heterothermy they reduce the set point of temperature regulation thus promoting heat loss instead of heat conservation, until this set point is attained (Geiser, 2011; Humphries et al., 2002; Mejias et al., 2022; Nespolo et al., 2021). Thus, communal nesting should be important at very low temperatures, as animals need to maintain heat to avoid tissue freezing. For instance, *Myotis sodalist* bats (8g), which hibernates at similar temperatures as in *D. gliroides* (i.e., between 2 and 8ºC, see Humphries et al., 2002; Nespolo et al., 2021), cluster together in hundreds of individuals. These animals, however, do not benefit from clustering during deep hibernation. The main energetic benefit of clustering is to reduce the cost of rewarming, since large groups reduce the thermal gradient for warm up the body (Boyles et al., 2008; a similar effect has been found in sugar gliders, see Nowack and Geiser, 2016)).

The sole study where the whole hibernation cycle was followed in monitos was performed in a coastal location with a mild climate (winter mean T_A_ ∼ 8ºC; Nespolo et al., 2022a). These authors found that clustering (and the use of hibernacula) is more frequent in the proportion of individuals that remain normothermic during the cold season, a common fact in this opportunistic hibernator (Bozinovic et al., 2004; Oda et al., 2019). This is coincident with the previous observation, indicating that heterothermic animals do not benefit from huddling if temperatures are not extremely low. To study how these behavioral strategies vary between lowland and highland populations remains as an open important question (some high-altitudinal populations of this marsupial has sub-zero T_A_s for most of the winter, see Fonturbel et al., 2022; Mejias et al., 2021).

### Biophysical models, energy budget and conservation

The use of biophysical models in physiological ecology dates back to Warren and Porter seminar paper (Porter and Gates, 1969; Tracy, 1972, see an update in Kearney and Porter, 2009), who defined the energy budget of animals as a balance equation of energy losses and gains in a variety of environments, type of organisms and thermal conditions. Later, a number of authors have applied them for predicting a variety of contexts (e.g., climatic impact on species range: Huey et al., 2012; Kearney et al., 2009; heat loss and behavioral thermoregulation in endotherms: Kenagy et al., 2002; McCafferty et al., 2011; reconstructing dinosaur physiology Seebacher, 2003), and recently, for predicting habitat suitability in mammalian conservation (McComb et al., 2021). This approach has the advantage of simplifying the experimental design, avoiding having to handle live animals in the laboratory, or to embark on costly fieldworks. Here we show that simple simulations could provide quantitative data for testing a specific hypothesis about the origin of sociality in a marsupial.

## Acknowledgements

This work was funded by ANID – Millennium Science Initiative Program – Center Code NCN2021-050 to RFN, and ANID PIA/BASAL FB0002 to FB and RFN.

## References

Altamirano, T. A., Honorato, M. T., Ibarra, J. T., de la Maza, M., de Zwaan, D. R., Bonacic, C. and Martin, K. (2019). Elevation has contrasting effects on avian and mammalian nest traits in the Andean temperate mountains. Austral Ecology 44, 691–701.

Altamirano, T. A., Ibarra, J. T., de la Maza, M., Navarrete, S. A. and Bonacic, C. (2015). Reproductive life-history variation in a secondary cavity-nester across an elevational gradient in Andean temperate ecosystems. Auk 132, 826–835.

Antinuchi, C. D. and Busch, C. (2001). Reproductive energetics and thermoregulatory status of nestlings in pampas mice Akodon azarae (Rodentia: Sigmodontinae). Physiological and Biochemical Zoology 74, 319–324.

Boratynski, J. S., Willis, C. K. R., Jefimow, M. and Wojciechowski, M. S. (2015). Huddling reduces evaporative water loss in torpid Natterer’s bats, Myotis nattereri. Comparative Biochemistry and Physiology a-Molecular & Integrative Physiology 179, 125–132.

Boyles, J. G., Storm, J. J. and Brack, V. (2008). Thermal benefits of clustering during hibernation: a field test of competing hypotheses on Myotis sodalis. Functional Ecology 22, 632–636.

Bozinovic, F., Rosenmann, M. and Ruiz, G. (1987). Transferencia de calor, convección y gradiente altitudinal. Archivos de Biolog-a y Medicina Experimentales 20, 85–88.

Bozinovic, F., Rosenmann, M. and Veloso, C. (1988). Termorregulación conductual en Phyllotis darwini (Rodentia:Cricetidae): efecto de la temperatura ambiente, uso de nidos y agrupamiento social sobre el gasto de energ-a. Revista Chilena De Historia Natural 61, 81–86.

Bozinovic, F., Ruiz, G. and Rosenmann, M. (2004). Energetics and torpor of a South American “living fossil”, the microbiotheriid Dromiciops gliroides. Journal of Comparative Physiology B-Biochemical Systemic and Environmental Physiology 174, 293–297.

Canals, M., Rosenmann, M. and Bozinovic, F. (1989). Energetics and geometry of huddling in small mammals. Journal of Theoretical Biology 141, 181–189.

Canals, M., Rosenmann, M. and Bozinovic, F. (1997). Geometrical aspects of the energetic effectiveness of huddling in small mammals. Acta Theriologica 42, 321–328.

Celis-Diez, J. L., Hetz, J., Marin-Vial, P. A., Fuster, G., Necochea, P., Vasquez, R. A., Jaksic, F. M. and Armesto, J. J. (2012). Population abundance, natural history, and habitat use by the arboreal marsupial Dromiciops gliroides in rural Chiloe Island, Chile. Journal of Mammalogy 93, 134–148.

Dausmann, K. H. and Glos, J. (2015). No energetic benefits from sociality in tropical hibernation. Functional Ecology 29, 498–505.

Ebensperger, L. and Bozinovic, F. (2000). Communal burrowing in the hystricognath rodent, Octodon degus:a benefit of sociality? Behavioral Ecology and Sociobiology 47, 365–369.

Ebensperger, L. and Labra, A. (2020). Comportamiento social de la fauna nativa de Chile: Ediciones UC.

Ebensperger, L. A. (2001). A review of the evolutionary causes of rodent group-living. Acta Theriologica 46, 115–144.

Ebensperger, L. A., Hurtado, M. J., Soto-Gamboa, M., Lacey, E. A. and Chang, A. T. (2004). Communal nesting and kinship in degus (Octodon degus). Naturwissenschaften 91, 391–395.

Fisher, D. O., Nuske, S., Green, S., Seddon, J. M. and McDonald, B. (2011). The evolution of sociality in small, carnivorous marsupials: the lek hypothesis revisited. Behavioral Ecology and Sociobiology 65, 593–605.

Fonturbel, F. E., Franco, L. M., Bozinovic, F., Quintero-Galvis, J. F., Mejias, C., Amico, G. C., Vázquez, M. S., Sabat, P., Sanchez-Hernandez, J. C., Watson, A. J. et al. (2022). The ecology and evolution of the Monito del monte, a relict species from the southern South America temperate forests. Ecology and Evolution ece3.8645.

Franco, M., Contreras, C., Cortes, P., Chappell, M. A., Soto-Gamboa, M. and Nespolo, R. F. (2012). Aerobic power, huddling and the efficiency of torpor in the South American marsupial, Dromiciops gliroides. Biology Open 1, 1178–1184.

Franco, M., Quijano, A. and Soto-Gamboa, M. (2011). Communal nesting, activity patterns, and population characteristics in the near-threatened monito del monte, Dromiciops gliroides. Journal of Mammalogy 92, 994–1004.

Geiser, F. (2011). Hibernation: Endotherms. eLS, 1–10.

Gilbert, C., McCafferty, D., Le Maho, Y., Martrette, J. M., Giroud, S., Blanc, S. and Ancel, A. (2010). One for all and all for one: the energetic benefits of huddling in endotherms. Biological Reviews 85, 545–569.

Gurovich, Y., Stannard, H. J. and Old, J. M. (2015). The presence of the marsupial Dromiciops gliroides in Parque Nacional Los Alerces, Chubut, Southern Argentina, after the synchronous maturation and flowering of native bamboo and subsequent rodent irruption. Revista Chilena De Historia Natural 88, Article 17.

Heenan, C. B. and Seymour, R. S. (2011). Structural support, not insulation, is the primary driver for avian cup-shaped nest design. Proceedings of the Royal Society B-Biological Sciences 278, 2924–2929.

Honorato, M. T., Altamirano, T. A., Ibarra, J. T., De la Maza, M., Bonacic, C. and Martin, K. (2016). Composition and preferences regarding nest materials by cavity-nesting vertebrates in the Andean temperate forest of Chile. Bosque 37, 485–492.

Huey, R. B., Kearney, M. R., Krockenberger, A., Holtum, J. A. M., Jess, M. and Williams, S. E. (2012). Predicting organismal vulnerability to climate warming: roles of behaviour, physiology and adaptation. Philosophical Transactions of the Royal Society B-Biological Sciences 367, 1665–1679.

Humphries, M. M., Thomas, D. W. and Speakman, J. R. (2002). Climate-mediated energetic constraints on the distribution of hibernating mammals. Nature 418, 313–316.

Kauffman, A. S., Paul, M. J., Butler, M. P. and Zucker, I. (2003). Huddling, locomotor, and nest-building behaviors of furred and furless Siberian hamsters. Physiology & Behavior 79, 247–256.

Kearney, M. and Porter, W. (2009). Mechanistic niche modelling: combining physiological and spatial data to predict species’ ranges. Ecology Letters 12, 334–350.

Kearney, M., Porter, W. P., Williams, C., Ritchie, S. and Hoffmann, A. A. (2009). Integrating biophysical models and evolutionary theory to predict climatic impacts on species’ ranges: the dengue mosquito Aedes aegypti in Australia. Functional Ecology 23, 528–538.

Kenagy, G. J., Nespolo, R. F., Vasquez, R. A. and Bozinovic, F. (2002). Daily and seasonal limits of time and temperature to activity of degus. Revista Chilena De Historia Natural 75, 567–581.

Lubbe, N., Czenze, Z. J., Noakes, M. J. and McKechnie, A. E. (2018). The energetic significance of communal roosting and insulated roost nests in a small arid-zone passerine. Ostrich 89, 347–354.

Madikiza, Z. J. K. and San, E. D. (2020). Patterns of nest box sharing in woodland dormice (Graphiurus murinus): Evidence for intra-sexual tolerance and communal nesting. Behavioural Processes 177.

McCafferty, D. J., Gilbert, C., Paterson, W., Pomeroy, P. P., Thompson, D., Currie, J. I. and Ancel, A. (2011). Estimating metabolic heat loss in birds and mammals by combining infrared thermography with biophysical modelling. Comparative Biochemistry and Physiology a-Molecular & Integrative Physiology 158, 337–345.

McComb, L. B., Lentini, P. E., Harley, D. K. P., Lumsden, L. F., Eyre, A. C. and Briscoe, N. J. (2021). Climate and behaviour influence thermal suitability of artificial hollows for a critically endangered mammal. Animal Conservation.

Mejias, C., Castro-Pastene, C. A., Carrasco, H., Quintero-Galvis, J. F., Soto-Gamboa, M., Bozinovic, F. and Nespolo, R. F. (2021). Natural history of the relict marsupial Monito del Monte at the most extreme altitudinal and latitudinal location. Ecosphere 12, 1–15.

Mejias, C. J. N.,, Sabat, P., Franco, L. M., Bozinovic, F. and Nespolo, R. F. (2022). Body composition and energy savings by hibernation in the South American marsupial Dromiciops gliroides: a field study applying quantitative magnetic resonance. Physiological and Biochemical Zoology (in press), 1–10.

Nespolo, R. F., Fonturbel, F. E., Mejias, C., Contreras, R., Gutierrez, P., Oda, E., Sabat, P., Hambly, C., Speakman, J. R. and Bozinovic, F. (2022a). A Mesocosm Experiment in Ecological Physiology: The Modulation of Energy Budget in a Hibernating Marsupial under Chronic Caloric Restriction. Physiological and Biochemical Zoology 95, 66–81.

Nespolo, R. F., Mejias, C., Espinoza, A., Quintero-Galvis, J. F., Rezende, E. L., Fonturbel, F. E. and Bozinovic, F. (2021). Heterothermy as the norm, homeothermy as the exception: variable torpor patterns in the South American marsupial monito del monte (Dromiciops gliroides). Frontiers in Physiology 12, Article 682394.

Nespolo, R. F., Opazo, J. C., Rosenmann, M. and Bozinovic, F. (1999). Thermal acclimation, maximum metabolic rate and nonshivering thermogenesis in Phyllotis xanthopygus (Rodentia) inhabiting the andean range. Journal of Mammalogy 80, 742–748.

Nespolo, R. F., Saenz-Agudelo, P., Mejias, C., Quintero-Galvis, J. F., Peña, I., Sabat, P., Sanchez-Hernandez, J. C. and Gurovich, Y. (2022b). The physiological ecology of the enigmatic colocolo opossum, the monito del monte (genus Dromiciops) and its role as a bioindicator of the broadleaf biome. In Marsupials as Bioindicators, vol. (in press) (ed. M. L. Larramendy). London: Royal Society of Chemistry.

Nicol, S. C. and Andersen, N. A. (2007). Cooling rates and body temperature regulation of hibernating echidnas (Tachyglossus aculeatus). Journal of Experimental Biology 210, 586–592.

Nowack, J. and Geiser, F. (2016). Friends with benefits: the role of huddling in mixed groups of torpid and normothermic animals. Journal of Experimental Biology 219, 590–596.

Oda, E., Rodriguez-Gomez, G. B., Fonturbel, F. E., Soto-Gamboa, M. and Nespolo, R. F. (2019). Southernmost records of Dromiciops gliroides: extending its distribution beyond the Valdivian rainforest. Gayana 83, 145–149.

Olson, M. N., Bowman, J. and Burness, G. (2018). Social thermoregulation does not explain heterospecific nesting in North American flying squirrels. Biological Journal of the Linnean Society 123, 805–813.

Patil, V. P., Morrison, S. F., Karels, T. J. and Hik, D. S. (2013). Winter weather versus group thermoregulation: what determines survival in hibernating mammals? Oecologia 173, 139–149.

Porter, W. P. and Gates, D. M. (1969). Thermodynamic equilibria of animals with environment. Ecological Monographs 39, 227–244.

Rezende, E. L. and Bacigalupe, L. D. (2015). Thermoregulation in endotherms: physiological principles and ecological consequences. Journal of Comparative Physiology B-Biochemical Systemic and Environmental Physiology 185, 709–727.

Ruf, T. and Geiser, F. (2015). Daily torpor and hibernation in birds and mammals. Biological Reviews 90, 891–926.

Russell, E. M. (1984). Social-behavior and social-organization of marsupials. Mammal Review 14, 101–154.

Scholander, P. (1955). Evolution of climatic adaptations in homeotherms. Evolution 9, 15–26.

Seebacher, F. (2003). Dinosaur body temperatures: the occurrence of endothermy and ectothermy. Paleobiology 29, 105–122.

Selonen, V., Hanski, I. K. and Wistbacka, R. (2014). Communal nesting is explained by subsequent mating rather than kinship or thermoregulation in the Siberian flying squirrel. Behavioral Ecology and Sociobiology 68, 971–980.

Sharbaugh, S. M. (2001). Seasonal acclimatization to extreme climatic conditions by black-capped chickadees (Poecile atricapilla) in interior Alaska (64 degrees N). Physiological and Biochemical Zoology 74, 568–575.

Statistica. (2006). StatSoft INC. STATISTICA (data analysis software system) version 6.1.

Tracy, C. (1972). Newotn’s law: its applicability for expression heat losses from homeotherms. Bioscience 22, 656–659.

Vazquez, M. S., Ibarra, J. T. and Altamirano, T. A. (2020). Austral Opossum adjusts to life in second-growth forests by nesting outside cavities. Austral Ecology 45, 1179–1182.

Vickery, W. L. and Millar, J. S. (1984). The energetics of huddling by endotherms. Oikos 43, 88–93.

Vogt, F. D. and Lynch, R. (1982). Influence of ambient temperature, nest availability, huddling, and daily torpor on energy expenditure in the white-footed mouse Peromyscus leucopus. Physiological Zoology 55, 56–63.

Willis, C. K. R. and Brigham, R. M. (2007). Social thermoregulation exerts more influence than microclimate on forest roost preferences by a cavity-dwelling bat. Behavioral Ecology and Sociobiology 62, 97–108.

Withers, P. C. and Jarvis, J. U. M. (1980). The effect of huddling on thermoregulation and oxygen consumption for the naked mole-rat. Comparative Biochemistry and Physiology A 66A, 215–219.

Wojciechowski, M. S., Jefimow, M. and Pinshow, B. (2011). Heterothermy, and the energetic consequences of huddling in small migrating passerine birds. Integrative and Comparative Biology 51, 409–418.

